# Female and male microglia are not different in the dentate gyrus of postnatal day 10 mice

**DOI:** 10.1101/2021.08.11.456021

**Authors:** Danielle Guez-Barber, Lorianna M. Colon, Dana Raphael, Max A. Wragan, Sanghee Yun, Amelia J. Eisch

## Abstract

Microglia, the resident immune cells of the brain, support normal brain function and the brain’s response to disease and injury. The hippocampal dentate gyrus (DG) is an area of microglial study due to its central role in many behavioral and cognitive functions. Interestingly, microglia and related cells are distinct in female vs. male rodents, even in early life. Indeed, postnatal day (P)-dependent sex differences in number, density, and morphology of microglia have been reported in certain hippocampal subregions at specific ages. However, sex differences in the DG have not yet been assessed at P10, a rodent developmental time point translationally relevant to human term gestation. To address this knowledge gap, Iba1+ cells in the DG (which are enriched in the Hilus and Molecular Layer) in female and male C57BL/6J mice were analyzed for their number (via stereology) and density (via stereology and via sampling). Next, Iba1+ cells were classified into published morphology categories. Finally, the percent of Iba1+ cells in each morphology category was multiplied by total cell number to generate a total number of Iba1+ cells in each morphological category. Results show no sex difference in Iba1+ cell number, density, or morphology in the P10 Hilus or Molecular Layer. The lack of sex difference in Iba1+ cells in P10 DG using commonly-employed methodologies (sampling, stereology, morphology classification) provides a baseline from which to interpret microglia changes seen after injury.

**HIGHLIGHTS:**

- Sex differences in Iba1+ cells were assessed in the dentate gyrus (DG) of P10 mice
- Iba1+ cells were assessed in DG microglia-rich subregions: Hilus and Molecular Layer
- Both stereology and sampling approaches were used to quantify Iba1+ cells
- No sex difference found in Iba1+ cell number, density, or morphology in P10 mice DG

## 1. INTRODUCTION

Microglia are brain immune cells that play critical roles in normal brain function and development [1–4] and support the brain’s response to disease and injury [5,6]. Microglia are also distinct in female vs. male rodents, even in the early postnatal period [7–10]. Sex differences in microglia are particularly notable in the rodent hippocampus—a brain region critical for diverse brain functions such as memory and mood regulation—as early life sex differences have been reported in hippocampal microglia number, density, and morphology [11–13]. Of note, several papers have reported the sex difference in hippocampal microglia “flips” over the course of early life, but the directionality of this flip varies depending on the approach and species studied. For example, female rats have less microglia in the hippocampal CA1 subregion than males at postnatal day (P) 4, but more at P30 [11]. In relative contrast, female mice have more volume of cells that express Iba1 (a protein expression in microglia) in CA1 than males at P8 and less at P28 with no sex difference in volume seen at P15 [12]. These and other studies use a range of approaches to quantify microglia, from stereology to sampling, underscoring the utility of collecting several microglia measures (number, density, morphology) to enable comparison with the existing literature [14–16]. A full understanding of baseline sex differences in microglia, particularly in the hippocampus, is fundamental to neuroscience investigations that probe the role of microglia in perinatal brain injury and later life cognition and behavioral function.

Notably absent from this literature on sex difference and hippocampal microglia is investigation at P10, a time point in mouse development that neuroanatomically correlates with full-term human gestation [17]. Given the interest in studying and manipulating microglia in the perinatal period [13,18], it is surprising that to-date there are no studies on sex differences in microglia in the mouse P10 hippocampus. Another knowledge gap in the literature is the paucity of data on microglia sex differences in the main “gateway” to the hippocampus, the dentate gyrus (DG). In rodents and humans, a substantial portion of DG cellular development and circuit maturation occurs in the postnatal period, and these processes—such as postnatal neurogenesis—are sex-dependent [19]. Given the potent role of microglia in postnatal DG neurogenesis [4,20,21], it is important to know if the number or morphology of DG microglia differ by sex in early life. Studies on sex differences in DG microglia in the rat exist [11,13], but surprisingly none have been performed at P10; no work has examined sex differences in DG microglia in the mouse.

To address these knowledge gaps, here we counted the number of Iba1+ microglia in the DG of female and male P10 mice. To match quantification approaches in the literature and potentially address conflicting results [11,13,22–26], we used stereology to collect total number and calculate the density of Iba1+ microglia and also assess density using a sampling approach. We examined Iba1+ cell morphology, as this is also commonly reported in the literature. This information was collected from two DG subregions where Iba1+ microglia are most dense in the P10 mouse: the Molecular layer (Mol) and Hilus. The resulting data are important as they examine sex differences in Iba1+ microglia in the mouse hippocampus at P10, an understudied but important postnatal timepoint.

## 2. METHODS

### 2.1 Animals

Pregnant female C57BL/6J mice were shipped from Jackson Laboratory on embryonic day 14 (E14). Mouse pups were kept in the same cage as the dam and their littermates following birth. Pups from 4 litters from 4 distinct dams were used for this P10 analysis (pups: 11 females, 9 males); each litter included both female and male pups. All mice were cared for in compliance with protocols approved by the Institutional Animal Care and Use Committee at CHOP and guidelines established by the NIH’s *Guide for the Care and Use of Laboratory Animals*. Our scientific reporting adheres to the ARRIVE 2.0 guidelines [27]. Details in **Supplementary Material 1.1 (S1.1)**.

### 2.2 Brain collection and tissue preparation

Mice were sexed at P10 prior to brain collection based on anogenital distance and gonadal appearance [28]. Brains were fixed for 72 hours (h) at room temperature (RT) in 4% paraformaldehyde (PFA), cryoprotected (30% sucrose, 0.1 M PBS), and sectioned (40μm, coronal through the anterior hippocampus on a microtome (**S1.2**).

### 2.3 Immunohistochemistry (IHC)

Slide-mounted chromogenic IHC was performed [29,30] with rabbit-anti-Iba1 (**S1.3**).

### 2.4 Stereological counting

Unbiased stereology (optical fractionator probe) was used to manually count Iba1+ cells in DG subregions. The Mol and Hilus were delineated using a low-power 5x objective (Zeiss AxioImager M2 microscope) on DG sections (**Fig.1A**) beginning at -1.35mm from bregma through the next 2400μm, and using a section sampling fraction of 1/6 [30] (**Fig.1B, S1.4**).

**Figure 1.**
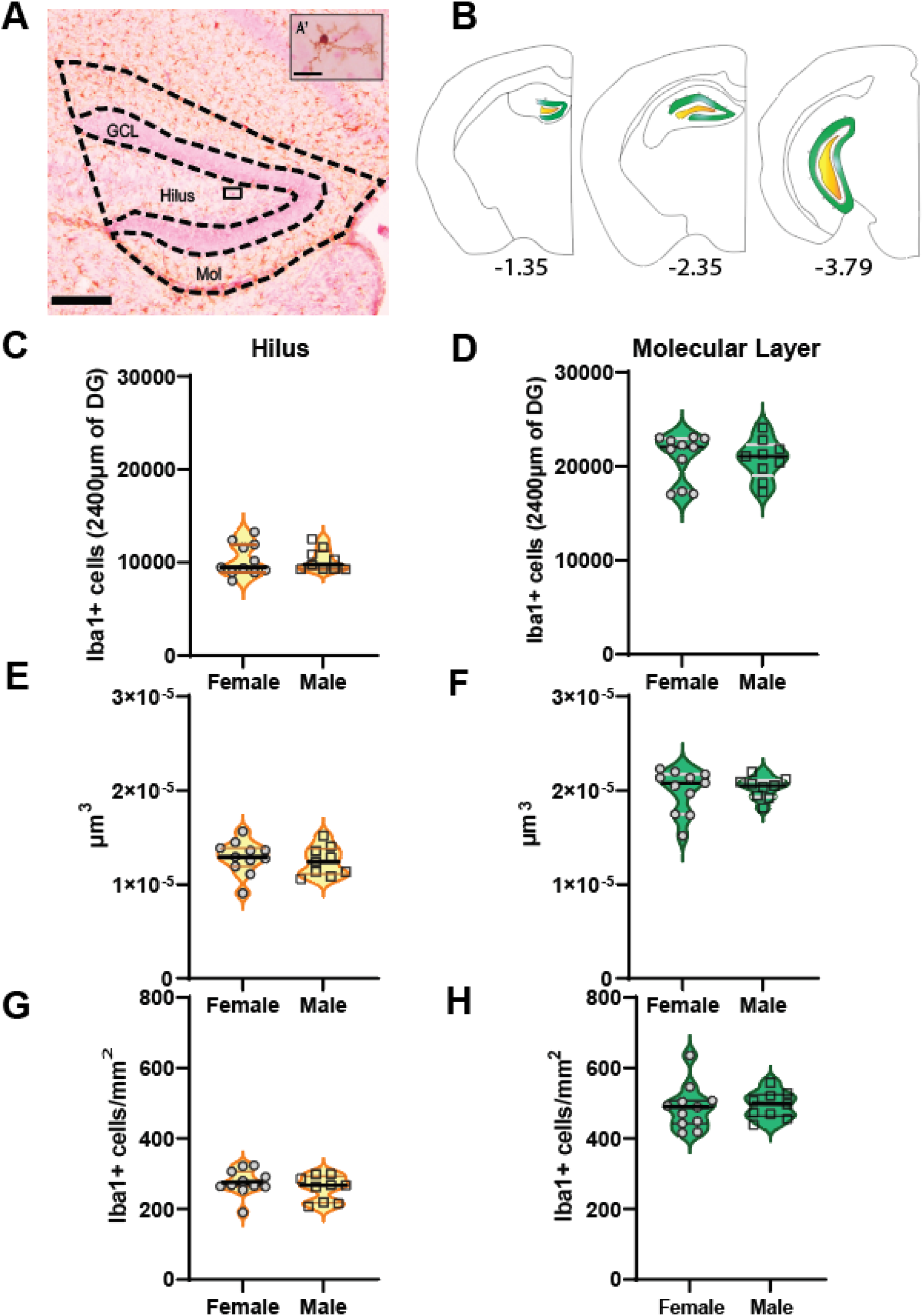
Stereology: No sex difference in Iba1+ cell number, density, or volume in P10 mouse dentate gyrus (DG). **(A)** Photomicrograph of a P10 mouse DG coronal brain section immunostained with anti-Iba1 antibody (FastRed counterstain). Dotted lines are DG subregions (Hilus, Molecular Layer [Mol]) in which Iba1+ cell number was assessed by the optical fractionator probe. The Granule Cell Layer (GCL) is also shown, but GCL Iba1+ cells were extremely rare and not quantified. Scale bar=200μm. Black Hilus rectangle enlarged in **(A’)** shows an Iba1+ Hilus cell with Thin morphology. Scale bar=25μm. **(B)** Stereotaxic atlas plates of coronal sections from Bregma -1.35 to -3.79 showing the Hilus (gold) and Mol (green) where Iba1+ cells were counted. Total Iba1+ cell number was quantified in the left and right hemispheres of P10 male and female mice within the anterior 2400μm of the DG, which included up to 10 hippocampal brain sections/mouse. There was no sex difference in Iba1+ cell **(C-D)** number, **(E-F)** volume, or **(G-H)** density in the Hilus **(C, E, G)** or Mol **(D, F, H)** of P10 mice. N=11 females, 9 males.

### 2.5 Stereological cell density

To determine the density of Iba1+ cells, volume of each region was determined via Cavalieri estimator probe. The total number of Iba1+ microglia for each mouse divided by the total volume of the region resulted in density (**S1.5**).

### 2.6 Sampling approach to density determination

Iba1+ microglia in photomicrographs of the Hilus and Mol were manually counted (**Fig.2A-B**). For each mouse, a total of four 400x images were collected (2 anterior DG regions from each hemisphere at -2.05 mm from bregma)[24,26](**S1.6**).

**Figure 2.**
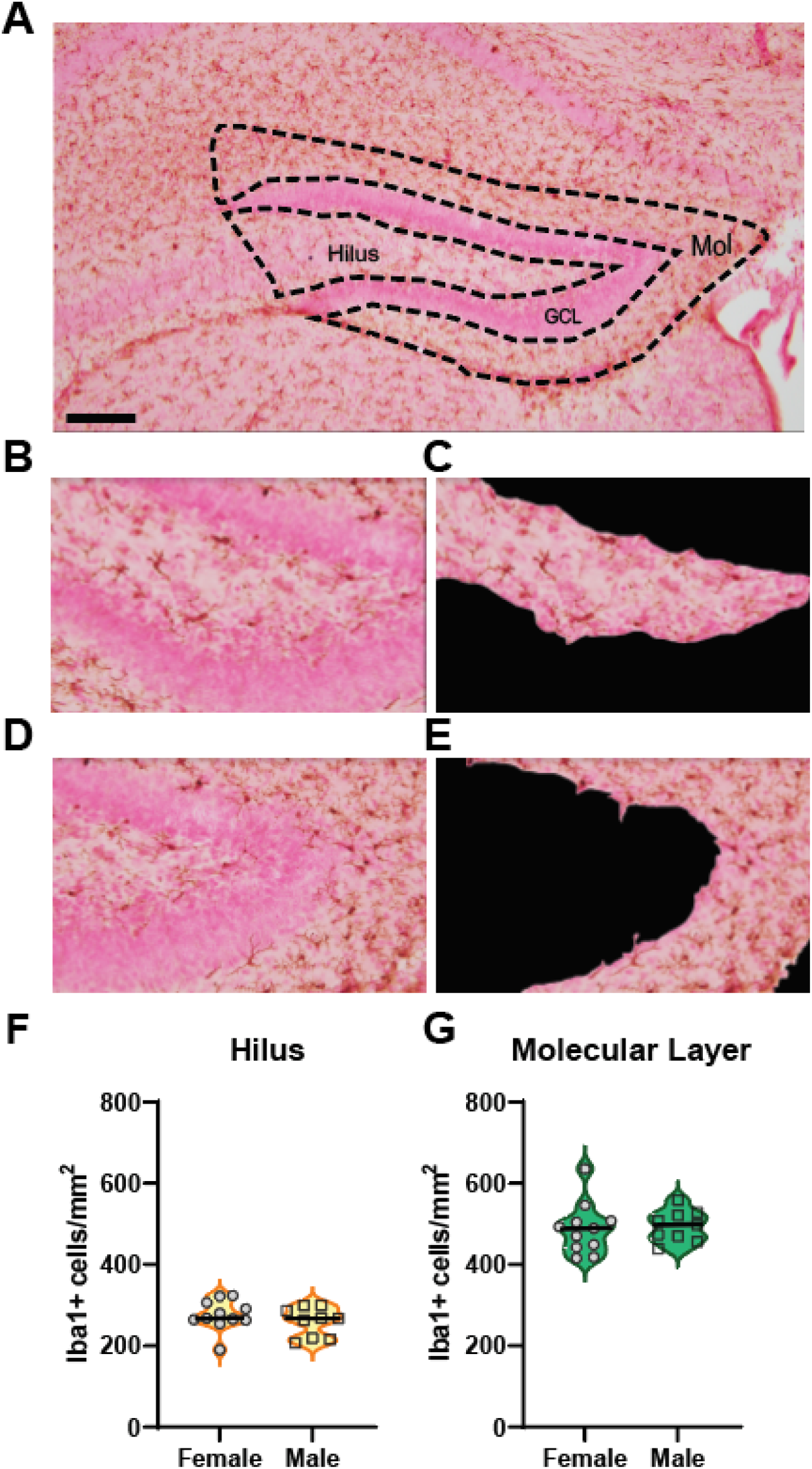
Sampling approach: No sex difference in Iba1+ cell density in P10 mice Hilus and Molecular Layer (Mol). **(A)** Photomicrograph of P10 anterior DG. Dotted lines indicate Hilus and Mol subregions. Scale bar=200um. **(B-E)** A region of interest (ROI) drawn around the Hilus **(B, D)** and Mol **(C, E)** in each photomicrograph enabled these Iba1+ cell-rich DG subregions to be assessed for Iba1+ cells while the granule cell layer (GCL) was shaded and not assessed. **(F-G)** There was no sex difference in the density of Iba1+ cells in the Hilus **(F)** or Mol **(G)**. N=11 females, 9 males.

### 2.7 Image collection and morphology analysis

Each Iba1+ microglia examined via sampling was placed into one of four previously-published morphology categories (Round/Amoeboid, Stout, Thick, Thin; **Fig. 3A-E**) based on the soma shape and processes [11,13,31–33](**S1.7**).

**Figure 3.**
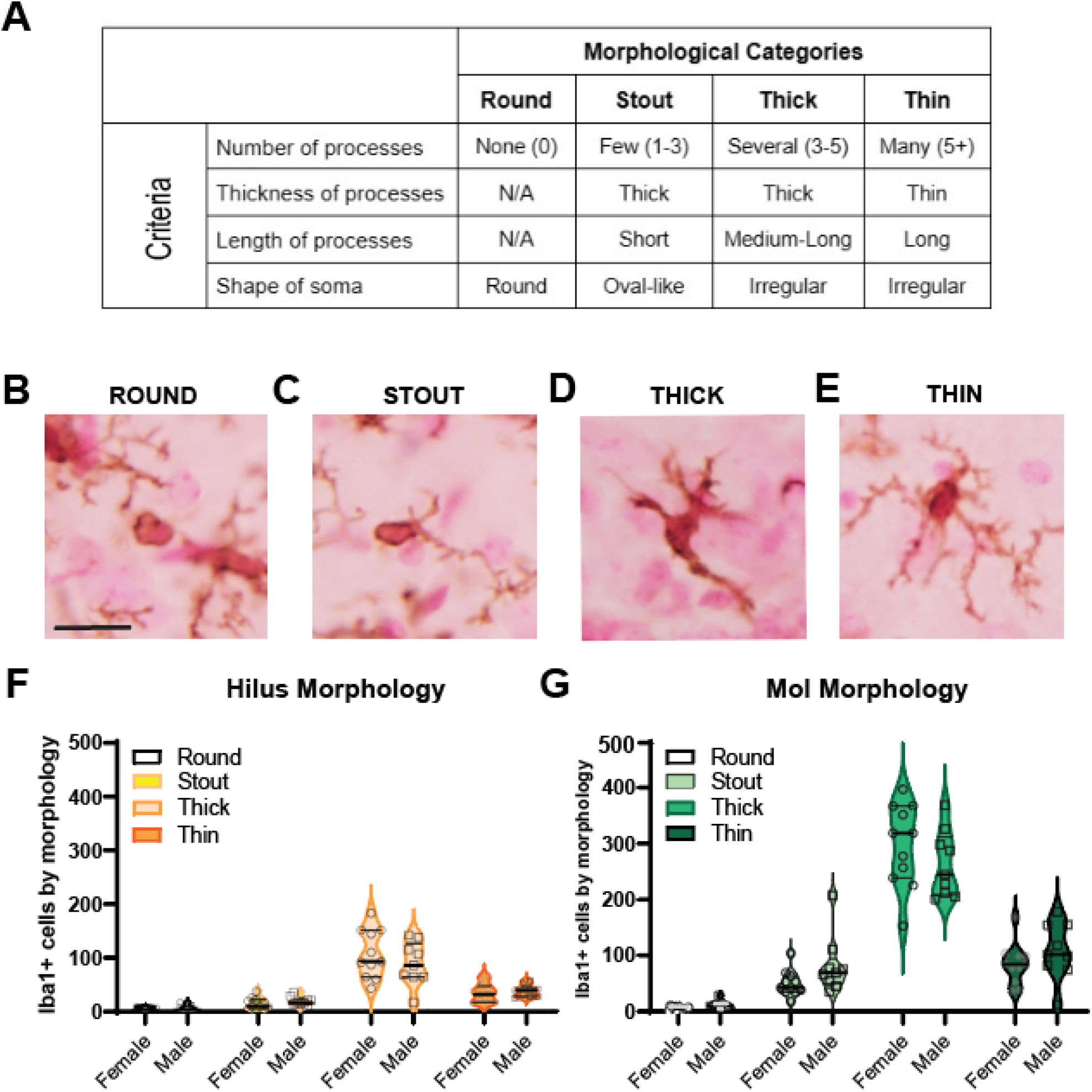
No sex difference in total number of Iba1+ cells by morphology in mouse P10 Hilus and Molecular Layer (Mol). **(A-E)** Criteria used for Iba1+ cell categorization into one of four morphologies **(A)**: **(B)** Round, **(C)** Stout, **(D)** Thick, or **(E)** Thin. **(F-G)** Iba1+ calculated cell number/morphology. Iba1+ Thick and Thin cells were most prevalent, while Iba1+ Round and Stout cells were far more rare or absent. Scale bar=25um, applies to **(B-D)**. N=11 females, 9 males.

### 2.8 Data analysis and statistics

Sex of the mice was masked until data collection was complete. Data were analyzed via Prism 9 (Graphpad; **S1.8**).

## 3. RESULTS

### 3.1 Stereology: No sex difference in Iba1+ microglia **number** in P10 DG subregions

Unpaired t-test of stereology data revealed no sex difference in total number in either the Hilus [*t*(18)=0.067; p>0.05, η^2^=0.0002,**Fig.1C**] or Mol [*t*(18)=0.154; p>0.05, η^2^=0.001,**Fig.1D**].

### 3.2 Stereology: No sex difference in Iba1+ microglia **density** in and total **volume** of P10 DG subregions

Unpaired t-test on Cavalieri data revealed no sex difference in total volume of the Hilus [*t*(18)=0.5438; p>0.05, η^2^=0.016,**Fig.1E**] or Mol [*t*(18)= 0.3150; p>0.05, η^2^=0.005,**Fig.1F**]. Unpaired t-test on the calculated density data revealed no sex difference in density of Iba1+ microglia in either the Hilus [*t*(18)=0.5271; p>0.05, η^2^=0.02,**Fig.1G**) or Mol [*t*(18)= 0.3402; p>0.05, η^2^=0.006,**Fig.1H**].

### 3.3 Sampling approach: No sex difference in Iba1+ microglia **density** in P10 DG

Unpaired t-test revealed no sex difference in cell density in either the Hilus [*t*(18)=1.012; p>0.05, η^2^=0.05,**Fig.2F**] or Mol [*t*(18)=0.2589; p>0.05, η^2^=0.004,**Fig.2G**].

### 3.4 No sex difference Iba1+ cell **morphology** in the P10 DG

Morphological analysis of the DG revealed four types of microglia commonly-seen in the literature [8,11,34]: Round, Stout, Thick, and Thin. A two-way ANOVA (Sex X Morphology) revealed no sex-dependent effects of morphology categorization in either the Hilus [F(3,72)=1.032; p>0.05,**Fig.3F**] or Mol [F(3,72)=2.162; p>0.05,**Fig. 3G**]. There was a main effect of Morphology in both the Hilus [F(3,72)=148.2; p<0.001,**Fig.3F)** and Mol [F(3,72)=319.3; p<0.001,**Fig. 3G**] such that in each subregion, Round and Stout cells were far more rare than Thick and Thin cells. This is consistent with few Round DG microglia reported at other earlylife timepoints [11].

## 4. DISCUSSION

Employing two common techniques (stereology and sampling [11,13,22–26]) we find no sex difference in Iba1+ cell number, density, or percent of Iba1+ cells in each morphology category in P10 DG subregions. The lack of sex-dependent difference in Iba1+ cell number or density in the P10 DG is interesting in regard to prior work. In rat DG, female and male microglia differ in early development: at P4 when most microglia have an immature morphology, males have more Round/Amoeboid microglia than females, but by P30 when most microglia have mature morphology, females have more Thick microglia than males [11]. That study examined many timepoints (ED17 to P60), but did not evaluate timepoints between P4 and P30 when this transition in rat was proposed to occur. Our work here in mice shows no sex-dependent difference in DG P10 Iba1+ cell number, density, or morphology, suggesting P10 might be a reasonable timepoint to assess for when this “flip” occurs. In fact, a study in mice showed females have more Iba1 volume vs. males at P8 and less vs. males at P28 [12]. It is notable that they found no sex difference in Iba1 volume P15; this aligns with our data showing no sex difference in Iba1+ cell number at P10. That work in mice concluded that hippocampal microglial development occurs earlier in females than males [12]. Another group using transcriptomics also concluded that microglia development is earlier in females than males [35]. This latter work is important to keep in mind when considering the implications of our present negative data; even when there are no sex differences in cell number, as we show here, there may be sex differences in gene expression patterns and other important phenotypes [13,36].

Our morphology data are similarly interesting, particularly in that most DG P10 Iba1+ cells are Thick or Thin (and few are Round/Amoeboid or Stout) in both sexes. These data were collected from immersion fixed brains; one concern may be Round cells are debris that would be removed if we exsanguinated. However, studies that use intracardial perfusion find Round microglia in the young and adult brain [8,11,34,37–39]. Also, the small number of Round Iba1+ microglia we report in the P10 DG is consistent with other early life timepoints [11]. Collectively this raises the question: why are most DG P10 Iba1+ cells Thick or Thin, with few being Round or Stout? This may reflect microglia maturity, with Round microglia (no processes) less mature than ramified Thin microglia [11]. Another possibility comes from the adult rodent, where similar morphology categories are used as a gradient to reflect the degree of microglia “activation”. However, reevaluation of the nomenclature used to describe and define microglia [1] suggest “activation” or “resting” do not adequately capture the dynamic nature of microglia, and that microglia should be considered in the context of the age, sex, and health/disease state.

There are limitations of this study. First, we presume that Iba1+ cells are microglia. However, Iba1 is expressed in several brain cells: brain-resident yolk-sac derived microglia, infiltrating peripheral monocyte-derived macrophages, and border-associated macrophages [40]. Although some of the Iba1+ cells analyzed here may be one of the latter two categories, prior research suggests this is a very low percentage, particularly in the uninjured brain [25,41]. While definitive assessment of the Iba1+ cells analyzed in our work as microglia vs. infiltrating macrophages requires co-labeling with distinguishing markers [25,42], we believe it is appropriate to consider that most of these Iba1+ cells are microglia. Second, the morphology categorizations were based on a single plane of focus when ideally this is done with multiple focal planes, 3D analysis, or even Sholl analysis of arborization.

Nevertheless, the same approach was used for both female and male brain sections by blinded analysis, so this would not have biased our work in a sex-specific manner. Third, in this work we only analyzed the hippocampal DG; future work should also examine microglia number, density and morphology in the CA1 and CA3 regions of the hippocampus. Fourth, while we examined several litters and both sexes were included from each litter, the small number of mice per litter precluded meaningful statistical analysis of litter effects. Finally, the pups used for this study came from timed pregnant dams who were delivered to our facility during pregnancy. It remains to be seen if pups from dams bred in-house will have similar differences in morphology classification as presented here.

In sum, here we show that there is no difference in Iba1+ microglia number, density, or morphology between female and male mice in the P10 DG. Because P10 is a rodent developmental time point translationally relevant to human term gestation [17], this fills a key knowledge gap. While all future studies will require similar control groups, our demonstration of no sex difference in Iba1+ cells in P10 DG provides a baseline from which to interpret microglial changes seen after injury, which often do not consider baseline microglia values.

## 5. Funding and acknowledgements

DGB is supported by the Division of Neurology at the Children’s Hospital of Philadelphia (CHOP), “Remapping the Clinical Neurosciences through Translation and Innovation Training” (2T32NS091008; co-PIs: Jensen, Aguirre), several avenues of support from the CHOP Research Institute (the Supplemental Support for Clinicians to Pursue Research Training in Neuroscience, a Foerderer Award for Excellence, a K-Readiness Faculty Pilot Program Award), the Alavi-Dabiri Postdoctoral Fellowship Award from CHOP’s Intellectual and Developmental Disabilities Research Center, and the Child Neurologist Career Development Program CNCDP-K12 (NS098482, PI: Schlaggar). LMC is supported by the CHOP Postdoctoral Diversity Fellowship. HMP and MW were supported by the University of Pennsylvania’s (Penn) Center for Undergraduate Research and Fellowships (CURF). DR was also supported by CURF as well as by funds from being named a Penn Department of Neuroscience Scholar. SY is supported by a 2019 NARSAD Young Investigator Grant from the Brain and Behavior Research Foundation, a 2020 University of Pennsylvania Undergraduate Research Foundation grant, an NIH R01 grant (NS088555-05, PI: AM Stowe), a 2021 NASA HERO grant (80NSSC21K0814), and a Foerderer Award for Excellence from CHOP. AJE and this work are also supported by Bridge Funding from the CHOP Research Institute, an NIH R01 grant (NS088555-05, PI: AM Stowe), and a research grant from the Peter F. McManus Charitable Trust. We are indebted to Sharon Xie for biostatistical guidance, F. Chris Bennett, Graham Peet and Mariko Bennett for microglia imaging advice and discussions, Margaret McCarthy for discussions on microglia and sex differences in early life, and Kelly Jordan-Sciutto’s group for permission to use their imaging set-up in the initial stage of this project. Thanks to many Eisch Lab members and colleagues for helpful discussions and guidance, in particular Lyles R. Clark and Fred C. Kiffer, and to Ruthie E. Wittenberg for input and editing. We thank Haley M. Phillips and Kira Lu for preliminary help with early data analysis.

## Data Availability Statement

An electronic copy of all experimental data will be made available upon reasonable written request.

