## Supplemental Materials and Methods for "Female and male microglia are not different in the dentate gyrus of postnatal day 10 mice"

### S1. Supplemental Materials and Methods

#### *S1.1* Animals

Pregnant multiparous adult female C57BL/6J mice (~12 weeks old) were shipped from Jackson Laboratory (stock #000664) on embryonic day 14 (E14). Dams were individually-housed in a HEPA-filtered, closed airflow vivarium rack (Lab Products Inc., Enviro-Gard™ III) under a normal 12:12-hour light-dark cycle (06:00 on, 18:00 off) with ad libitum access to food (standard rodent chow, Lab Diets 5015 #0001328) and water. The AAALAC-accredited facility at the Children's Hospital of Philadelphia (CHOP) Research Institute is temperature- (22°C) and humidity-controlled (30-70%). Mice were weighed once prior to tissue collection at P10.

#### *S1.2* Brain collection and tissue preparation

Mice were euthanized on P10 via live decapitation using sterilized, surgical scissors in compliance with NIH guidelines for euthanasia of rodent neonates. Once extracted from the skull, meninges were removed and brains were fixed for 72 hours (h) at room temperature (RT) in 4% paraformaldehyde (PFA). Following two PFA changes (one/day), brains were cryoprotected via placement in 30% sucrose in 0.1 M PBS. Forty µm coronal sections at 200 µm intervals through the anterior hippocampus were collected on a freezing microtome (Leica SM 2000 R) and stored in 1xPBS with 0.01% sodium azide at 4°C until processing for immunohistochemistry (IHC).

#### *S1.3* Immunohistochemistry (IHC)

As previously described [1,2], brain sections were mounted onto charged slides (Thermo Fisher Scientific, #12-550-15) and left to dry for 1-2 h prior to IHC. Sections underwent antigen retrieval (0.01 M citric acid, pH 6.0, 100°C, 15 min) followed by RT washes in 1xPBS; all subsequent steps also occurred at RT. Non-specific protein binding was blocked via incubation with 3% Normal Donkey Serum (NDS) and 0.3% TritonX-100 in 1xPBS for 60 minutes (min). Following pretreatment and blocking steps, sections were incubated overnight with rabbit-anti-Iba1 (1:500; Wako Chemicals, #019-19741) in a carrier of 3% NDS and 0.3% Tween-20 in 1xPBS. Eighteen hours later, sections were rinsed with 1xPBS prior to incubation with biotinylated donkey-anti-rabbit IgG (1:200; Jackson ImmunoResearch Laboratories, #711-065-152) in 1.5% NDS. After washing with 1xPBS, endogenous peroxidase activity was inhibited via incubation with 0.3% hydrogen peroxide in 1xPBS for 30 min. Following an additional set of 1xPBS washes, slides were incubated with an avidin-biotin complex (ABC Elite, Vector Laboratories, #PK-6100) for 90 min. Slides were washed with 1xPBS and immunoreactive cells were visualized via incubation with metal-enhanced diaminobenzidine (Thermo Fisher Scientific, #34065) for 11-12 min. Slides were counterstained via Fast Red (Vector Laboratories, #H3403), dehydrated in a series of increasing ethanol concentrations and Citrosolv, and coverslipped with DPX (Thermo Fisher Scientific, #50-980-370).

*S1.4* Stereological counting
To manually count Iba1+ cells in DG subregions, we used unbiased stereology via the optical fractionator probe (Stereo Investigator, MBF Bioscience, Williston, VT, USA). All stereology data were collected by a person blinded to sex of the mouse. The Mol and Hilus were delineated using a low-power 5x objective (Zeiss AxioImager M2 microscope) on DG sections (main text, **Fig. 1A**) beginning at -1.35mm from bregma through the next 2400μm, and using a section sampling fraction of 1/6 as explained in prior work [2]. To delineate the Hilus, a line was drawn from the tip of the infrapyramidal blade to the tip of the suprapyramidal blade of the DG and around the inner border of the GCL. To delineate the Mol, following the trajectory of the Hilus, a line was extended up from the edge of the suprapyramidal blade to the hippocampal fissure, across the outer border of the GCL, and along the border of the infrapyramidal blade. Iba1+ microglia were manually counted (40x objective) with a counting frame of 60x60μm (Hilus) and 50x50μm (Mol), a dissector height of 10μm, and a sampling grid of 125x125μm based on a Schaeffer coefficient error of ≤0.01 [3,4]. Section thickness was determined via z-plane analysis at each counting site. Total cell number was based on the number weighted section thickness. All Iba1+ microglia bodies that came into focus within the counting frame or crossing acceptance lines without touching rejection lines [5] and met our criteria for size (~10μm), color (even staining), and shape (round, stout, thick or thin) were manually counted. The total estimated population using number weighted section thickness was generated by the optical fractionator probe; this value represents the total number Iba1+ cells within the first 2400μm of the DG. In cases where a section of tissue was damaged or folded in the region of interest, the missing section tool in the optical fractionator probe was utilized and the estimate adjusted by missing section fraction value (Stereo Investigator) was used to provide an unbiased measure of the total number of Iba1+ microglia. Of the 399 sections analyzed (200 Mol and 199 Hilus sections), the missing section tool was utilized on only 13 sections.

*S1.5* Stereological cell density
To determine the density of Iba1+ cells, volume of each region was determined via Cavalieri estimator probe (Stereo Investigator; section thickness 40μm, grid spacing 100μm), similar to prior work [6–8]. The total number of Iba1+ microglia for each mouse divided by the total volume of the region resulted in density.

*S1.6* Iba1+ cell density determination via sampling
Images from immunostained sections were captured in bright-field light microscopy using an Olympus BX51 microscope with a 40x/0.9NA objective and an Olympus DP74 camera (widescreen aspect ratio, 16:10). Guidelines for taking photomicrographic images of the DG subregions (Mol, Hilus; main text, **Fig. 2A-B**) were established to enable anatomical matching of subregions across hemispheres and mice and to reduce inter-person variation in taking photomicrographs. These guidelines were as follows: Mol: the outer Mol near the DG crest was positioned in the bottom and lateral corner of the image. Hilus: the DG crest was placed in the lateral and central screen such that the Hilus extended into the middle of the image. After photomicrographs were collected, a region of interest (ROI) was drawn on each image using FIJI by an observer blinded to the mouse's sex. This ROI ensured the appropriate regions would be analyzed (e.g. Mol) and other regions that may be in the image would be excluded (main text, **Fig. 2C-D**), and enabled quantification of total pixels within each ROI. For analysis, ROI areas (in pixels) were converted to $um^2$ based on 1 pixel=0.0231 $um^2$. Iba1+ cells were examined in each RO1. Criteria for inclusion as an Iba1+ microglia were a consistently-labeled soma easily distinguishable from the counterstain and the presence of an entire cross-sectional view through the soma in the photomicrograph focal plane. The photomicrographs for each subregion were annotated to indicate each Iba1+ microglia that was manually counted and its morphology

classification. This enabled a permanent record of the data collection and also allowed data collection validation as additional observers of the original, non-annotated photomicrographs could compare their results with the annotated versions. Iba1+ microglia count for one hemisphere for each of the DG subregions was divided by the area of the ROI to generate an Iba1+ cell density for that hemisphere (main text, **Fig. 2E-F**). Density values for each hemisphere were averaged such that each mouse was represented by one density value for each subregion.

*S1.7* Image collection and preparation for morphology analysis

After morphological categorization as previously published [9–13], the number of cells in each category was transformed into a percentage based on the total number of Iba1+ cells classified in the bilateral anterior hippocampus. A percentage for each of the four categories (Round, Stout, Thick, and Thin) was generated for each hemisphere in each subregion. The four percentages from hemisphere subregions were averaged with the other hemisphere subregion percentage, such that each mouse subregion was represented by four category percentages. The percentage of each morphological cell type in the Hilus and Mol were multiplied by the total number of cells generated via stereology to produce a single value for total number of cells by each morphological classification (main text,**Fig. 3E-F**).

*S1.8* Data analysis and statistics

Normality tests via group distribution were first assessed by the D'Agostino and Pearson test which revealed groups to be normally distributed. Normally-distributed Iba1+ total cell number via stereology and cell density measures via sampling and stereology for Hilus and Mol were next tested via t-test for an effect of sex. Normally-distributed Iba1+ morphology classification for each subregion was tested via 2-way ANOVA (main effects: Sex [Male, Female] X Morphology [Round, Stout, Thick, Thin]). Data are expressed as mean +/- SEM and $\alpha = 0.05$.
